# Macaque retina simulator

**DOI:** 10.64898/2026.03.09.710551

**Authors:** Simo Vanni, Francescangelo Vedele, Henri Hokkanen

## Abstract

The primate retina dissects visual scenes into multiple retinocortical streams. The most numerous retinal ganglion cell (GC) types, midget and parasol cells, are further divided into ON and OFF subtypes. These four GC populations have anatomical and physiological asymmetries, which are reflected in the spike trains received by downstream circuits. Computational models of the visual cortex, however, rarely take GC signal processing into account.

We have built a macaque retina simulator with the aim of providing biologically plausible spike trains for downstream visual cortex simulations. The simulator is based on realistic sampling density and receptive field size as a function of eccentricity, as well as on two distinct spatial and three temporal receptive field models. Starting from data from literature and earlier receptive field measurements, we synthetize distributions for receptive field parameters, from which the synthetic units are sampled.

The models are restricted for monocular and monochromatic stimuli and follow data from the temporal hemiretina which is more isotropic.

We show that the model patches conform to anatomical data not used in the reconstruction process and characterize the responses with respect to spatial and temporal contrast sensitivity functions. This simulator allows starting from a stimulus video and provides biologically plausible spike trains for the distinct unit types. This supports development of thalamocortical primate model systems of vision. In addition, it can provide a reference for more biophysical retina models. The independent parameters are housed in text files supporting reparameterization for particular macaque data or other primate species.

**Author summary:** Visual environment provides a rich source of information, and the visual system structure and function has been studied for decades in many species, including humans. The most complex data in mammalian species are processed in the cerebral cortex, but to date we are still missing a functioning model of cortical computations.

While the earlier anatomical and physiological data describe many details of the visual system, to understand the functional logic we need to numerically simulate the complex interactions within this system.

To pave the way for simulating visual cortex computations, we have developed a functioning model for macaque retina. The neuroinformatics comprises a review and re-digitized existing retina data from literature, as well as statistics of earlier macaque receptive field data. Finally, we provide software which brings the collected neuroinformatics to life and allows researchers to convert visual input into biologically feasible spike trains for simulation experiments of visual cortex.

## 1 Introduction

The retina dissects visual information into numerous parallel streams [1–3]. In the macaque, each eye has 1.4–1.8 × 10^6^ ganglion cells (GCs) distributed over an area of 670 mm^2^ [4,5]. Of the about 20 different GC types, roughly 90% of GCs send their axons to the lateral geniculate nucleus (LGN). The midget GCs comprise about 65% of all GCs, are sensitive to fine spatial structure and red-green contrast, and project to the parvocellular (P) layers of the LGN. The second most numerous is the parasol GC type (5 – 10%), which is sensitive to rapid changes in input and small changes in luminance contrast, and projects to the magnocellular (M) layers of the LGN. From the LGN, neurons in the P and M layers project to distinct layers of the primary visual cortex (V1, for a review, see [6]).

Functionally, it is advantageous to have a GC population specialized for high spatial and low temporal frequency content and another one specialized for low spatial and high temporal frequency content [7]. Behavioral measurements with targeted LGN lesions have shown that the P and M pathways are essential for conscious vision and that they carry distinct types of information [8–10]. Red-green color discrimination and recognition of fine patterns are dependent on the P pathway, while contrast sensitivity and recognition of stimuli with high temporal frequency depend on the M pathway.

The parasol and midget GC types are further divided into ON cells, sensitive for light increments, and OFF cells, which are more sensitive to light decrements. An analytical model showed that a system using ON units and OFF units needs less spikes for delivering the same information to downstream decoder, compared with a system using two ON units [11]. The OFF GCs have smaller dendritic fields and RFs and are more numerous than their ON counterparts [12,13], perhaps due to more dark contrast in natural scenes [14].

Our models are based on the large existing base of retina research. Vertebrate GC receptive fields were originally idealized as a linear circularly symmetric temporospatial filters [15], followed by a non-linear rectifying spike generation [16]. This model was named linear-nonlinear (LN) model, and it has been widely used as the model of the visual receptive field.

In the following decades, experimental data resulted in upending this early view. In the temporal domain, the LN model missed the saturating nature of contrast response, which was captured by a dynamic contrast gain control model, where contrast modifies temporal summation of the parasol cells [17,18]. Recently, part of the nonlinearity has been shown to emerge from rapid cone-mediated adaptation that allows photoreceptors to maintain optimal sensitivity despite frequent gaze shifts [19].

In the spatial domain, the circular receptive field structure was questioned when spike-triggered averaging measurements showed more asymmetry in the spatial structure of the receptive fields [12,20,21]. Later experiments revealed nonlinear summation in the parasol cell receptive field center [22,23], especially in natural viewing conditions, which was mechanistically explained by a shifting input signal gain at the bipolar to ganglion cell synapse [24].

Early modeling viewed GC spike generation as a Poisson process, but later data suggested a much more orderly temporal behavior, first in amphibians [25] and later in primates [26]. Moreover, photoreceptors were found to be very noisy. This noise is at a faster frequency than visual signaling and appears to result in correlated firing of dissimilar ganglion cell types [27]. As temporal dynamics have been suggested to guide the development of downstream circuits [28], we considered necessary to control the temporal signal and noise parameters.

The pathway splitting and nonlinearities highlight the complex processing of visual signals before they reach the cortex. Thus, any synthetic approach wishing to understand visual processing in the primate cortex should account for these retinal processes either together or as separate modeling hypotheses. To facilitate studies on cortical visual processing using spiking neural networks, we implemented two spatial and three temporal models of macaque GC responses with an option for including two types of cone noise as well as two distinct spike generation models. Compared to the classical LN model (Fixed temporal model), our other models emphasize either temporal (Dynamic temporal model) or spatial fidelity (Subunit temporal and VAE spatial model). In addition, distinct component combinations will support testing downstream sensitivity for particular retinal features.

## 2 Methods

We used data obtained from macaque monkeys and state explicitly when data comes from other *genera*.

### 2.1 Data sets

The two spatial models and the linear temporal model are based on a data set which comprised electrophysiological recordings in the peripheral macaque retina (low spatial resolution maps in [21]; obtained from E.J. Chichilnisky, personal communication, Aug 2019). The original Linear-Nonlinear-Poisson (LNP) model [20] consisted of a tonic drive component and a space-time separable spatiotemporal filter, with the spatial component having 13×13 pixels at 60 μm/pixel in retinal coordinates and the temporal component having 15 samples at 30 frames/second. Supplementary Table 1 presents the spatial and Supplementary Table 2 the temporal characteristics of this experimental data.

Descriptive or statistical distribution functions or variational autoencoder network were fitted to this data and these models were used for generating new model units.

All other data were extracted from other published studies. Original data and quantitative models were digitally reconstructed from figures and tables and these values are available in numpy npz or csv file formats in the code repository at macaqueretina/retina/literature_data.

### 2.2 Spatial receptive field layout

For initial positioning of units, the ganglion cell density data as a function of eccentricity were digitized from [4,29,30], the cone data from [31,32], and bipolar cell densities were estimated from cone densities using bipolar type specific bipolar to cone rations (Supplementary Table 3, from [33]).

The density of many GC types is inversely proportional to their dendritic field coverage, suggesting constant coverage factor [4,34]. Midget coverage factor is 1 [34–36]. It is likely that coverage factor is 1 for all our unit types [21,37]. At the moment, we are not able to reach constant coverage factor of 1 due to technical limitations in the way we optimize the receptive field positions, rotations and overlap.

The density of GC RFs cannot be estimated from GC density alone, since the GCs are displaced centrifugally near the fovea. Thus, we followed the estimate of GC sampling density at the fovea and actual density at the periphery, as in [38]. The GC sampling density for the most foveal 3 mm was estimated from cone density (digitized from [38]) with an exponentially decaying scaling factor (filled dots in Fig 1A; scaling factor = 2.4 * e^-1.1r^ + 0.9). The GC somas (open dots) align with their sampling density centrifugally from 3 mm. We fitted this data to a double exponential function

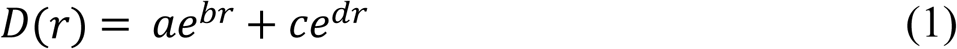

**Figure 1.**
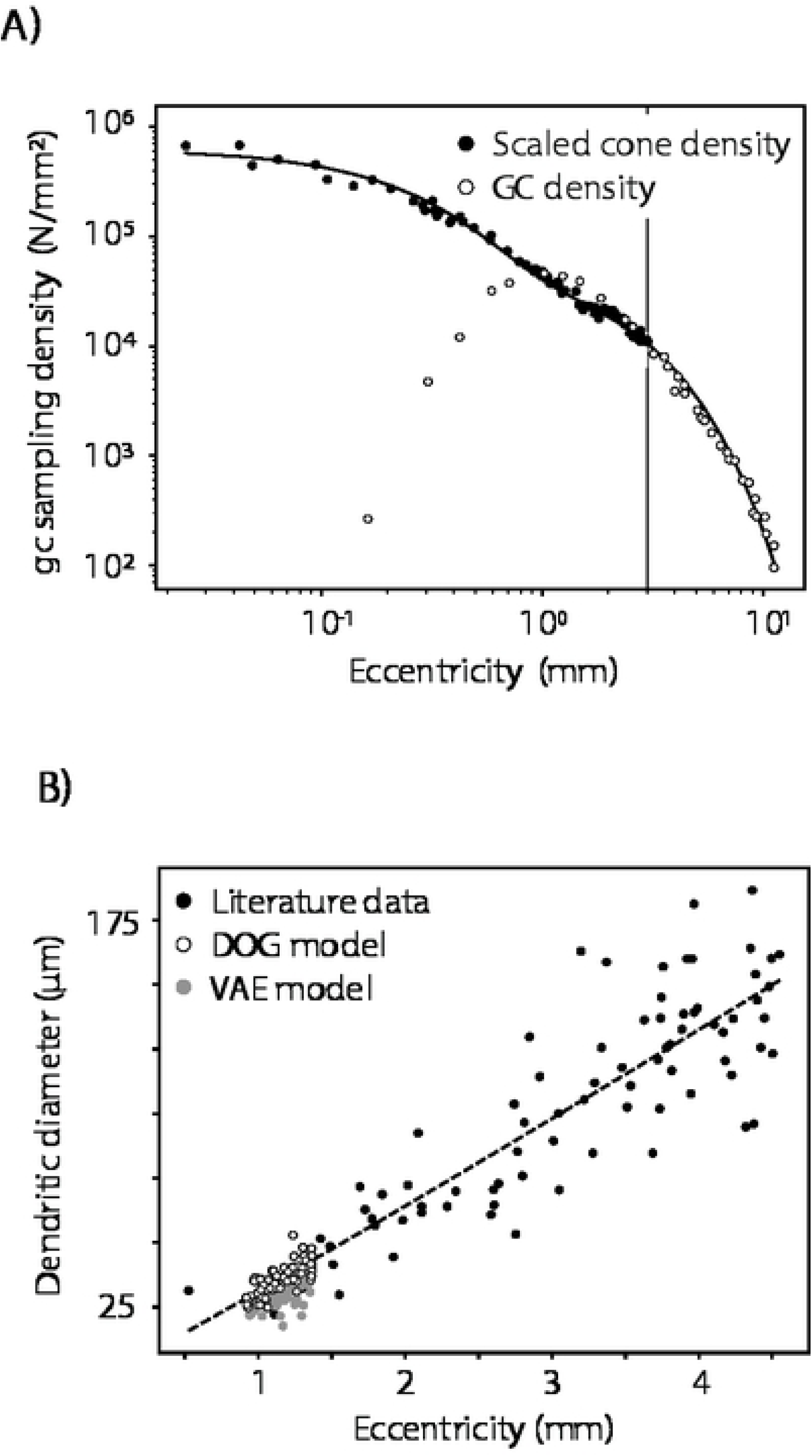
A) Ganglion cell sampling density of macaque temporal hemiretina, data digitized from [38]. GC sampling density for the most foveal 3 mm (filled dots) is estimated from cone density. The GC somas (open dots) are displaced from the fovea, but align with their sampling density centrifugally from 3 mm. The vertical line depicts the 3 mm eccentricity cutoff. B) Ganglion cell dendritic field diameter up to 20 deg eccentricity. Temporal hemiretina data (digitized from [4,29,30]). Example generated parafoveal DOG and VAE RF diameters are shown. The VAE model center diameter was estimated by fitting a descriptive difference-of-Gaussians model to the generated RF image.

where r is eccentricity (solid line in Fig 1A). The scaling parameters were *a* = 5.8 * 10^5^ and *c* 5.6 * 10^4^, and the corresponding decay rates were *b* = −4.2 and *d* −0.56. This descriptive model captured the GC sampling both at the fovea and in the periphery.

While midget and parasol cells form the majority of the parvo- and magnocellular-projecting subpopulations, respectively, there are other GC types projecting to these layers as well [39]. Earlier studies showed that there is an increase in the percentage of magnocellular-projecting cells towards the retinal periphery [30,40], but as this increase is rather modest in the temporal retina (6-10%), we did not take it into account. Thus, based on available estimates, we used the value 64% (of total GC density) for midgets and 8% for parasols, and we implemented a uniform midget-to-parasol proportion of 8:1 in our models.

Chichilnisky & Kalmar [12] reported that ON parasols have ∼20% larger receptive field (RF) diameters than the corresponding OFF cells. In humans, both midget and parasol cells show a similar asymmetry: the ON cell dendritic field diameters are 30-50% larger [13]. Based on these reports, we set the proportion of ON/OFF cell counts to 40%/60% for both midget and parasol GCs.

The GC dendritic field diameter, corresponding to their receptive field size, increases towards the periphery (Fig 1B and 2A). Parasol cell dendritic field diameter (digitized from [4,29,30]) up to 20 deg eccentricity has an almost linear power law regression slope of form 31.07 * E^1.05^ μm. For the full temporal hemiretina, the regression slope is 47.63 * E^0.76^ μm; this latter estimate was used for scaling the VAE model RFs according to eccentricity (Fig. 2A). For midget cells, the dendritic diameters close to the fovea do not shrink; instead, they show a tendency to increase, and these data were described with a quadratic function 11.6 – 6 * E + 2.8 * E^2^ μm. Figure 2B illustrates the key receptive field attributes for the VAE and DOG models.

**Figure 2.**
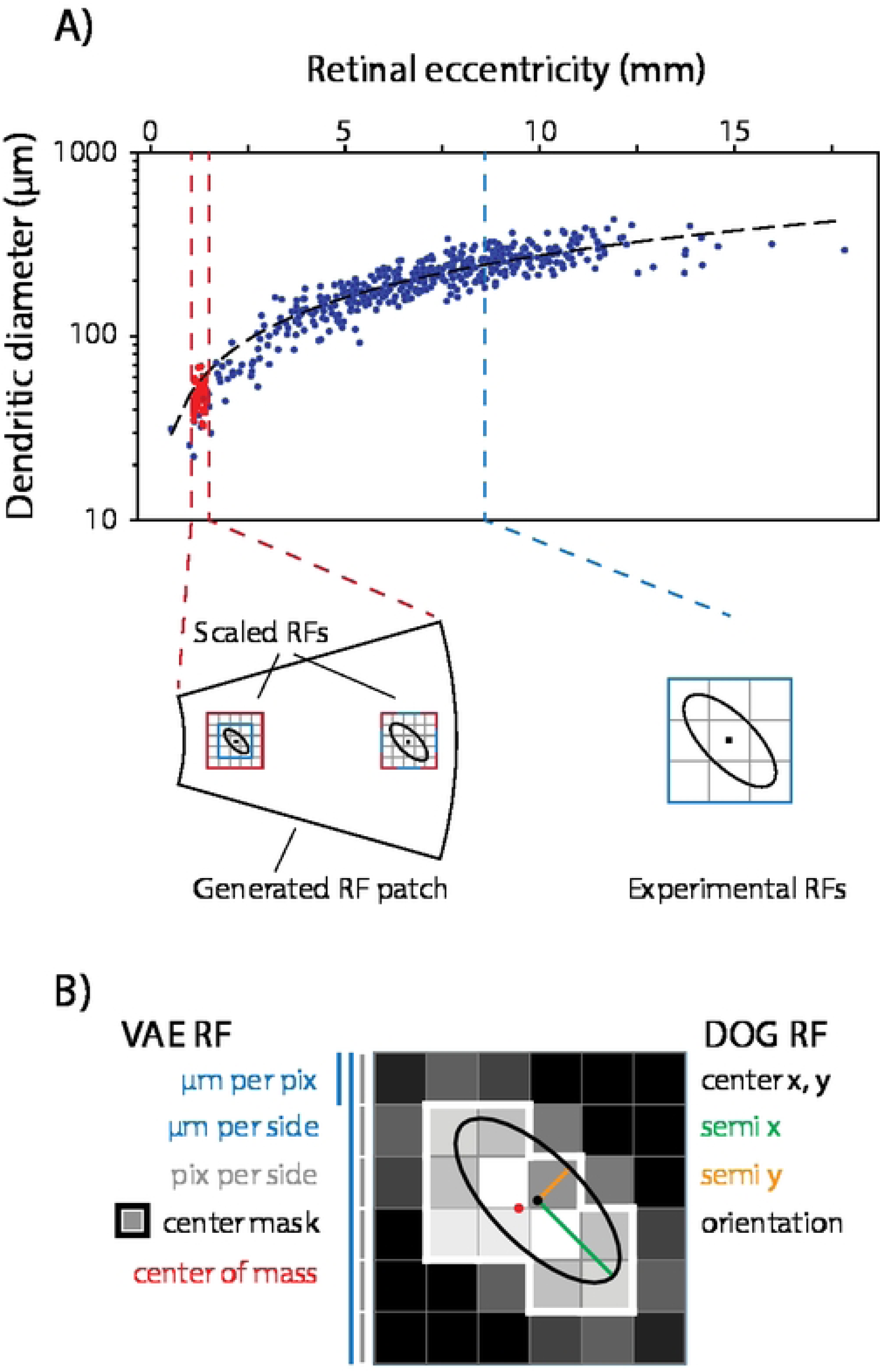
A) Receptive field spatial scaling as a function of eccentricity. Original spatial RF data [12,20,21] was from peripheral retina (blue dashed line). The corresponding dendritic field diameter data (blue dots, from [4,29,30]) provided a reference scale, and DOG RFs at other eccentricities were scaled following the descriptive power law regression slope (black dashed line). VAE models required additional resampling. First, we defined the minimum and maximum eccentricity for a synthetic retinal patch. Next, the pixel size (μm per pixel) is scaled according to the most foveal RF eccentricity, determining resolution, and the number of pixels (pix per side) is scaled according to the most peripheral RF eccentricity, determining the grid size. Finally, the RF images generated with the VAE model are scaled according to the regression slope and resampled to this new pixel grid. B) Key concepts for spatial receptive fields for VAE and DOG models. The center mask (pixels >10% of max value) is depicted with white outline.

### 2.3 Spatial models

Optical aberrations were fixed to 2 arcmin full width at half maximum, which corresponds to foveal data in humans [41]. Aberration was implemented with a 2D Gaussian smoothing function.

#### 2.3.1 Difference-of-Gaussian (DOG) models

We fitted the spatial filters to a difference of 2-dimensional elliptical Gaussian functions [15,42]:

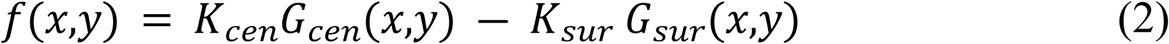

Here, *K_cen_* and *K_sur_* are the peak sensitivities of the center and surround, respectively. The Gaussians were parametrized by the center point, orientation, and SDs of the semimajor and semiminor axes (Fig. 2). The center and surround Gaussians were set to have the same center point, orientation, and aspect ratio. Allowing *K_cen_* to vary did not improve fits, so we fixed *K_cen_* at 1. We use the term “RF radius” to denote the geometric mean of the SDs along the major and minor axes, and “strength” to denote the volume under the Gaussian envelope. In the accompanying software, we have also implemented circularly symmetric and independent center and surround DOG models, but these were not analyzed further.

#### 2.3.2 Variational autoencoder (VAE) spatial model

To better capture the spatial statistics, we applied a variational autoencoder model to spatial RF data (Fig. 3A). The encoder started with a 32-dimensional latent space and 16 channels. The model employed a learning rate of 0.0005 and processed data in batches of 256 samples. Twenty percent of the data was allocated for both validation (10%) and testing (10%). The convolutional architecture employed two layers with kernel size 7 and stride 1, incorporating batch normalization. Original attempts to model the latent space with Gaussian distribution failed to provide other than spherical receptive fields, whereas a uniform distribution provided diverse RFs with a better match to data. Data augmentation was implemented with a 50% probability for both horizontal and vertical image flips, and the training dataset was expanded fourfold through augmentation procedures.

**Figure 3.**
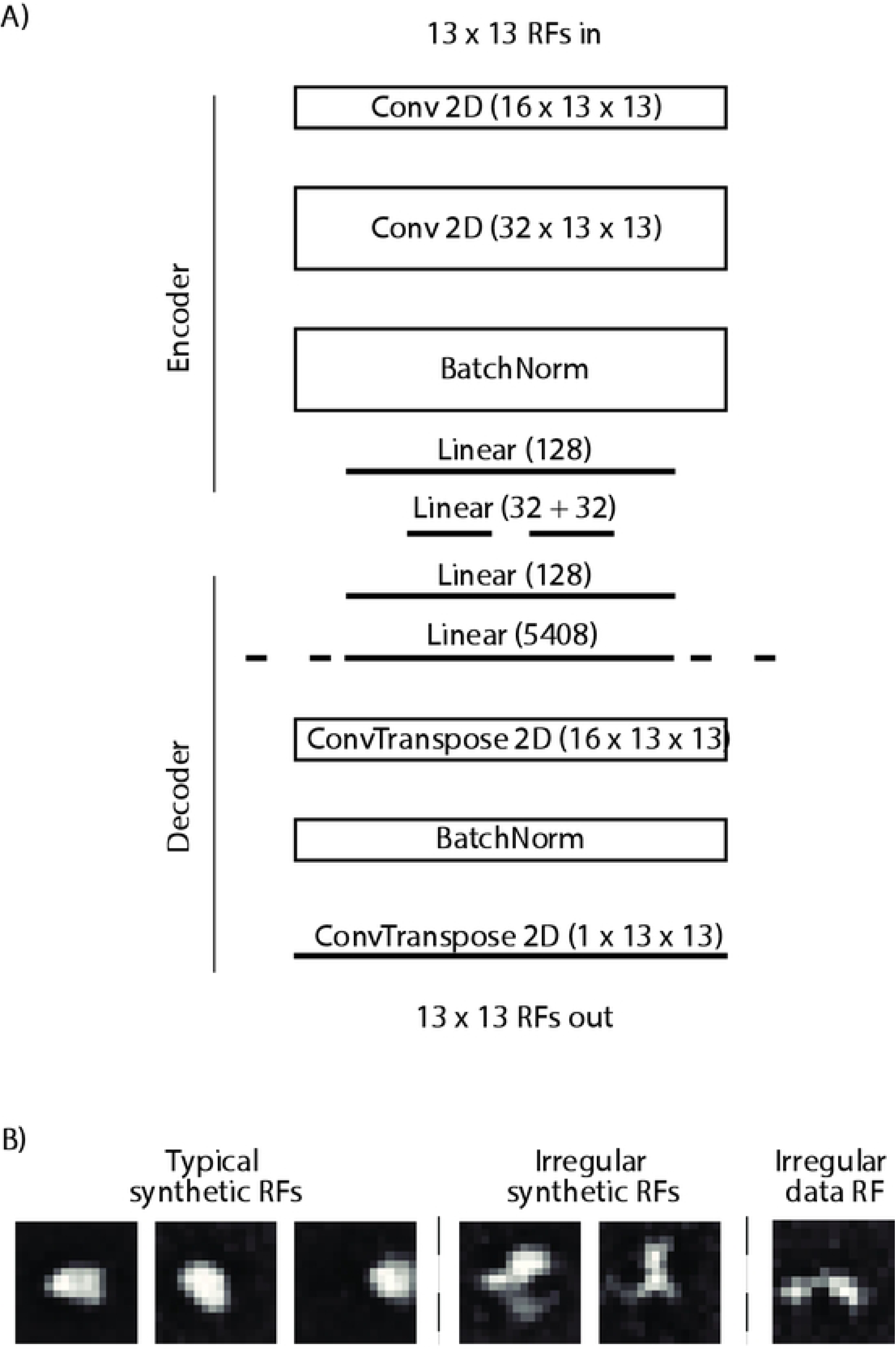
A) VAE model network structure. B) Example synthetic parasol ON receptive fields. Some asymmetric RFs emerged from the VAE (middle) and were found in the original data as well (right)

Figure 3B shows examples of regular and irregular synthetic spatial receptive fields, together with one example of irregular RF in the original primate data.

#### 2.3.3 Receptive field repulsion

Figure 4 shows the tiling of spatial RFs. Receptive field densities were estimated at 0.1 mm eccentricity intervals. First, receptive fields were compiled onto a regular hexagonal array. Next, spatial noise was added for the RF center positions. After selection of spatial model type (DOG vs VAE), receptive fields were tiled with random orientation. This resulted in an uneven sampling of the visual field (Fig. 4, top, parasol ON units). Next, we applied an iterative repulsion with receptive field rotations and translations. We used an objective of minimizing receptive field overlaps, resulting in more even distribution of the RF sampling (Fig 4, bottom). At this point, the amplitudes of the DOG and VAE RFs are disparate due to different generation processes.

**Figure 4.**
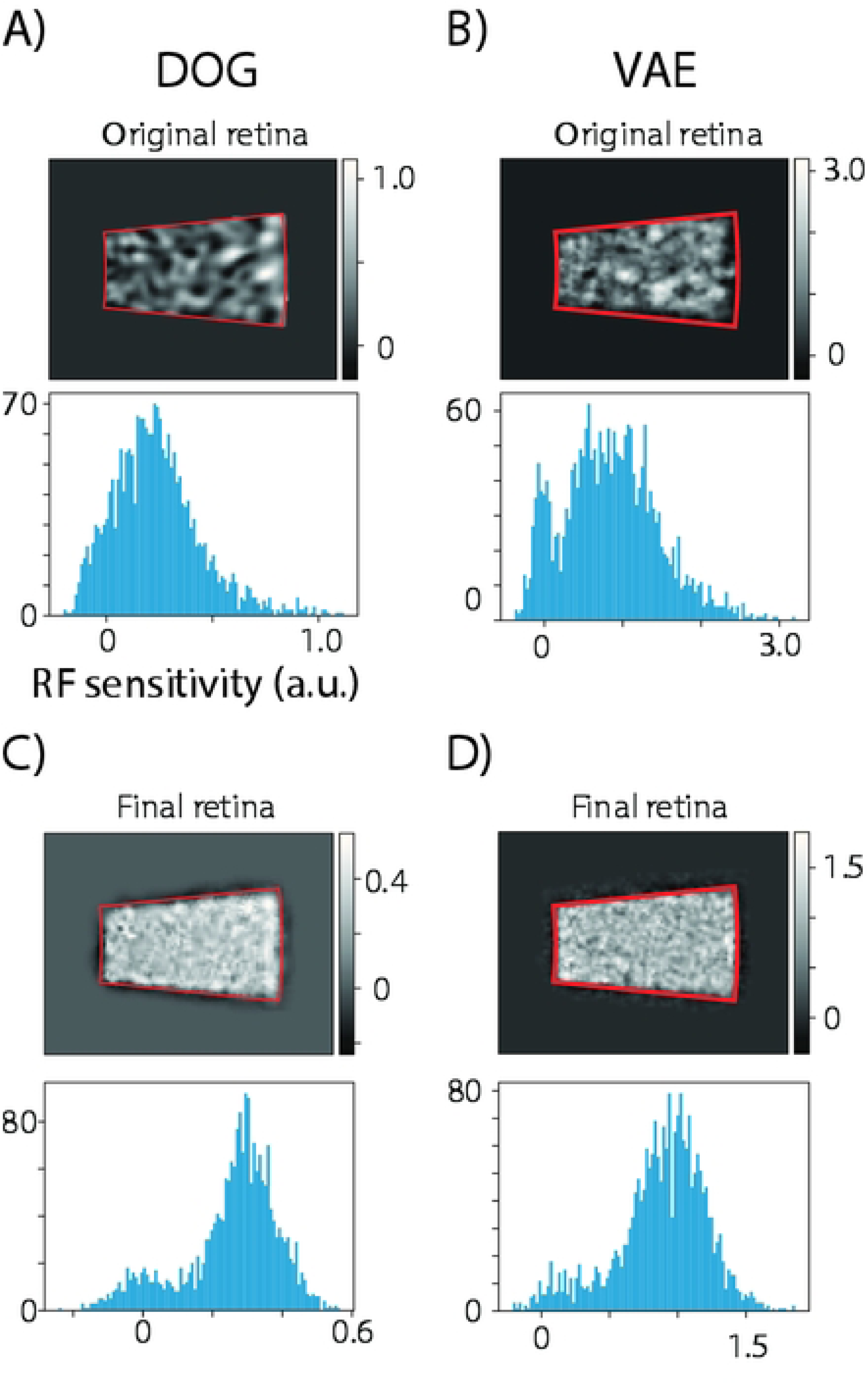
After tiling the RF center points, both the DOG (A) and VAE (B) RF centers sum into uneven distribution resulting in uneven sensitivity across the model retina. After repulsion, the final DOG (C) and VAE (D) RFs show narrower distribution and more even sensitivity.

### 2.4 Temporal models

We constructed three temporal models: a classical linear model (Fixed), a contrast gain control model (Dynamic), and a model combining fast cone adaptation with RF center subunit nonlinearity (Subunit). The parameters for the Fixed and Dynamic models describe a function from visual stimulus to ganglion cell response (end-to-end), whereas the Subunit model has a separate cone adaptation model followed by a bipolar subunit model. Thus, the Subunit model has no existing end-to-end parametrization from literature. The distinct temporal models were gain calibrated as explained in section 2.7.

#### 2.4.1 Fixed temporal model

A fixed temporal model assumes a linear transformation from image to generator potential. Thus, this transfer function is insensitive to changes in contrast or stimulus dynamics.

Figure 5 (black curves) depict Fixed model response for impulse (Fig 5A) and sustained (Fig 5B) luminance onset stimuli. The primate temporal response data points (N=15 at 30 frames per second) were fitted to a model consisting of the difference of low-pass filters [12]:

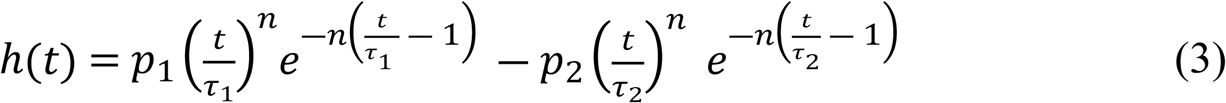

**Figure 5.**
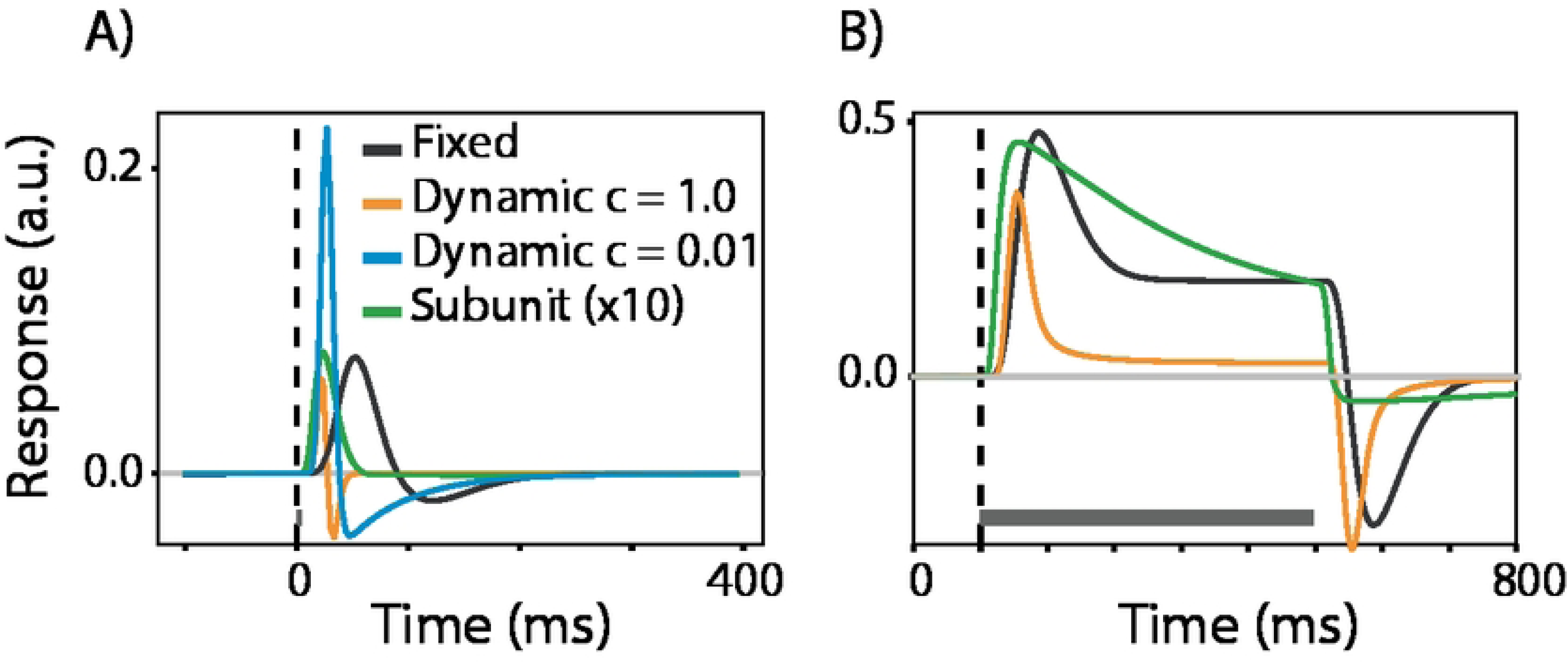
A) Generator potential (before nonlinearity and spike generation) impulse response function for the three temporal models. Note the distinct temporal kernel for low and high contrast (c) adaptation status for the Dynamic model. B) Corresponding traces for 500 ms luminance onset response.

Here, *t* is the time before present (in milliseconds) and *n, p_1_, p_2_, τ_1_, τ_2_* are free parameters.

Model parameter distributions were evaluated from the Chichilnisky data. New units were generated by sampling these distributions. It proved necessary to consider the correlations between the distributions in the sampling procedure, which limited the distribution models to multivariate Gaussian. Moreover, a hard sampling limit was set to 0-value, where appropriate, to avoid inverting the on-off polarity of the unit.

#### 2.4.2 Dynamic contrast gain control model

A dynamic gain control model [18] accounts for the response saturation at high contrasts in parasol cells, whereas the parametrization for midget cells is not contrast dependent. For reviews of physiological mechanisms of the gain control model, see [43] and [44].

Figure 5 shows an example of Dynamic model impulse response function for low (blue) and high (orange) contrast stimuli. Our Dynamic temporal model follows [18], where a linear low-pass stage precedes a dynamically-adjusted nonlinear high-pass stage.

Low-pass filter:

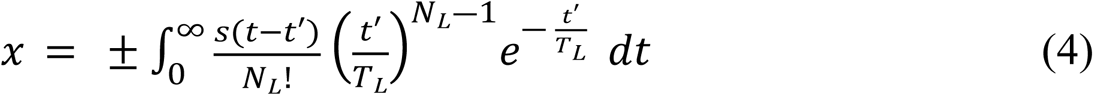

Here, *x* is the output vector from the linear low-pass stage. The input is *s(t)*, the signed Weber contrast at the receptive field center. The low-pass filter has *N_L_* stages of time constant *T_L_*. The ON-center cells have the initial + sign keeping the polarity of the response, while the OFF-center cells carry the – sign, reversing the polarity.

High-pass filter:

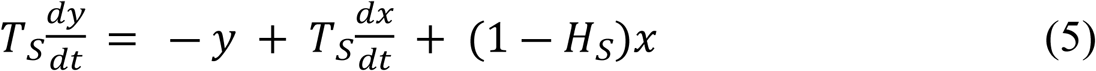

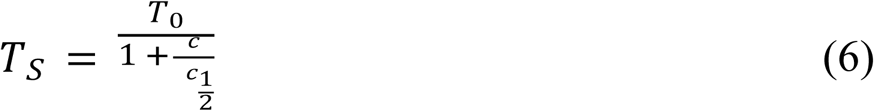

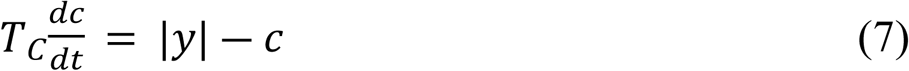

Here, *T_S_* is a dynamic time constant which depends on time-varying neural measure of contrast, denoted with *c*. The *c* is the low-pass filtered absolute magnitude of *y*, the output of the high-pass stage. *T_C_* is the time constant for this low-pass filter and 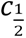 the contrast where *T_S_* is reduced by half.

The dynamic T_S_ is applied only to parasol units, whereas for midget units the T_S_ is constant due to their more linear dependence on contrast. The midget [45], and the parasol parameters [46] are presented in supplementary tables 1 and 2, respectively. Random samples were drawn from that data assuming triangular distribution.

#### 2.4.3 Subunit temporal model with fast cone adaptation

The cone adaptation model [47] with macaque parameters (from [19]) captures the rapid dynamical shift in cone response during one fixation.

The cone photoreceptor response *r* is modeled as a dynamical system driven by activation *y* and inactivation *z* inputs.

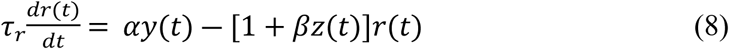

The τ_r_ is the time constant, and α and β are scaling constants. Light intensity *s* drives the *y* and *z* responses:

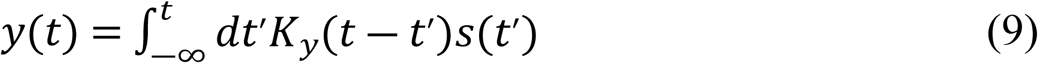

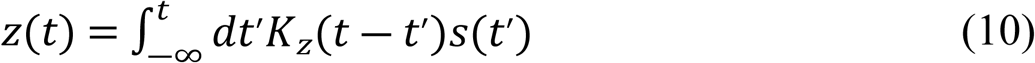

Where the K_y_ is the gamma distribution:

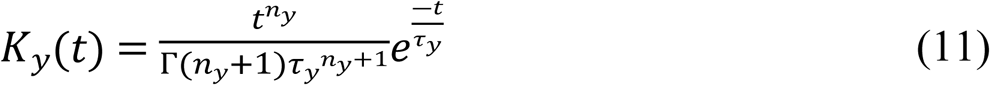

And the K_z_ is the weighted combination of K_y_ and an additional gamma filter:

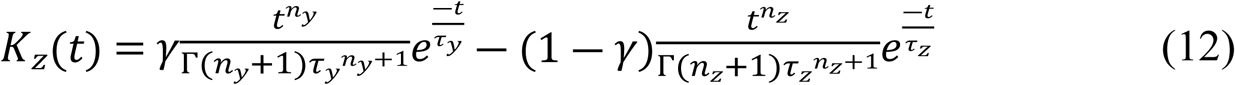

The normalized cone output c(t) corresponds to the relative synaptic vesicle release:

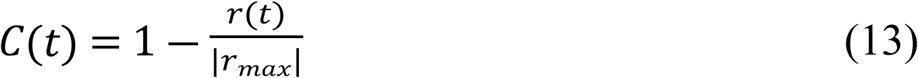

Where r_max_ is the maximum response amplitude (pA).

The cone vesicle release is maximal in darkness and decreases with higher retinal illumination. The cone-to-bipolar cell synapse operates with graded responses with sign preservation for the OFF bipolars and sign inversion for the ON bipolars:

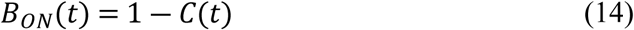

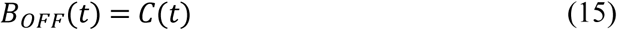

Next, we calculated bipolar contrast as:

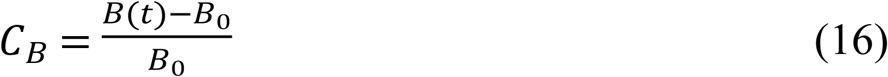

Where B_0_ is the bipolar input at the beginning of a sweep, typically mid gray background. C_B_ was our normalized measure of linear bipolar summation.

The subunit model emulates bipolar to ganglion cell synapse nonlinearity, which is guided by surround contrast [22–24]. If surround is weakly active or hyperpolarizes the center response, the center response becomes rectified. Instead, when the surround signal depolarizes the center, the center response becomes linearized. This relationship is distinct for ON and OFF parasols [12], and quantified with a rectification index RI (digited from [24]). RI near zero indicates a linear relationship between center input and output, whereas RI near one indicates a strong rectification. RI surround activation data was scaled to [-0.15,0.15] to match our maximal surround bipolar contrast, instead of the [-5,5], the excitatory synaptic conductance in [24].

We implemented the nonlinearity by scaling the negative ganglion cell input with 1-RI, a negative scaler. This scaler was applied at the bipolar unit level, separately for each timepoint.

An impulse response at photopic luminance (Fig 5A., green curve) shows a fast transient response followed by an almost nonexistent undershoot. Figure 5B depicts the response to 500ms full field light increment in ON units. The longer adaptation time results in a larger undershoot. Compared with the Fixed and Dynamic models, the Subunit model shows a clearly longer adaptation time constant. Differing from Fixed and Dynamic models, only the Subunit model dynamics change with cone illumination.

For subunit model we assume constant bipolar to cone ratio as function of eccentricity. The bipolar cell type-dependent parameters are from [33] (Supplementary Table 3) which report the bipolar densities at 6-7 mm eccentricity. Inputs to ON parasol ganglion cells come from Diffuse Bipolars, DB4 and DB5 [33,48], to OFF parasols from Diffuse Bipolars, DB2 and DB3 [49], to ON midgets from invaginating Midget Bipolars, IMB [50], and to OFF midgets from flat midget bipolars, FMB [50,51].

### 2.5 Spike generation

While a Poisson process is a good model of firing rate over sufficiently long intervals, GCs fire spikes more reliably than a Poisson process [26]. The spike generator’s precision can be improved by including post-spike feedback to model bursting, refractoriness and cell coupling [25,52]. We have implemented alternatives for a Poisson or a simple refractory spike generation model, following [25,26].

The temporal models output a generation potential which was first rectified:

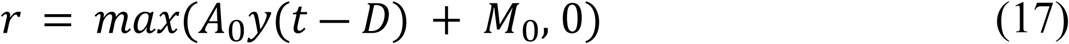

The firing rate *r* is the non-negative sum of the output high-pass stage, scaled by gain *A*_0_, and background firing rate *M*_0_. Output to the following processing stage has conduction delay *D*. Our M0 comprises the shared or Gaussian cone noise model (see below).

Next, this instantaneous firing rate passes through either a Poisson or Refractory spike generator model. The Refractory model [25,26] is of the form:

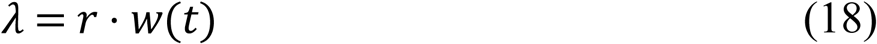

Where lambda is the instantaneous firing rate after refractory function w(t):

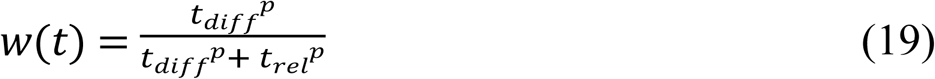

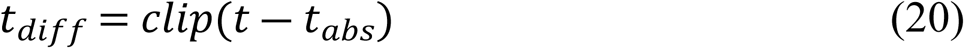

The absolute refractory period (t_abs_) was set to 1 ms, the relative refractory period (t_rel_) to 3 ms, and the exponent p was 4. Clipping sets negative numbers to zero.

### 2.6 Cone noise model

Ganglion cell spontaneous firing emerges primarily from cone noise, shared across the ganglion cell types [27]. The 4000 R* noise and response spectra were digitized from [53] and fit with a combination of cone response spectrum and double Lorentzian functions with two distinct (24 and 200-Hz) corner frequencies. The cone response spectrum covers the spontaneous photoisomerization noise, whereas the double Lorentzian function covers other phototransduction noise. This noise spectrum was used to generate spontaneous fluctuations in cone signals.

Spontaneous firing rates in midget and parasol cells ranges from 5 to over 30 Hz [54,55]; we fixed the cone noise to induce ∼25Hz baseline firing rates for the ON parasol and midget units, and ∼5Hz for the OFF parasol and midget units (Raunak Sinha, personal communication). In addition to this correlated noise, we implemented an alternative of unit-wise independent Gaussian noise with the same mean and SD. This allows exploration of possible functional significance of the correlated noise in downstream cortical computations.

### 2.7 Gain calibration between models

Numerical generation of different spatial and temporal RF models results in varying sensitivities to visual stimuli. To level this difference, we calibrated the GC generator potentials with a temporal contrast sensitivity experiment to drifting sinusoidal grating [56]. The parasol units used a 5 cpd sinusoidal grating for temporal contrast sensitivity calibration and 3.5% contrast for both calibrations, whereas the midget cells used 2 cpd grating and 11.4% contrast, correspondingly. We replicated this experiment at multiple gain values, resulting in a gain calibration table between distinct models.

Figure 6 shows the calibration procedure for DOG spatial and Fixed temporal model parasol ON units. First (Fig 6A), sinusoidal drifting grating was presented for 12 s at multiple frequencies and 10 different gain values. Resulting spike trains were binned, smoothed, and Fourier transformed. Next, we selected the F1 frequency providing the highest spike rate for most of the gain values. For the selected responses, a descriptive nonlinear function provides a good fit (Fig 6B). Next, we chose the gain producing 10-Hz response, in correspondence with rate used for most cells in [56].

**Figure 6.**
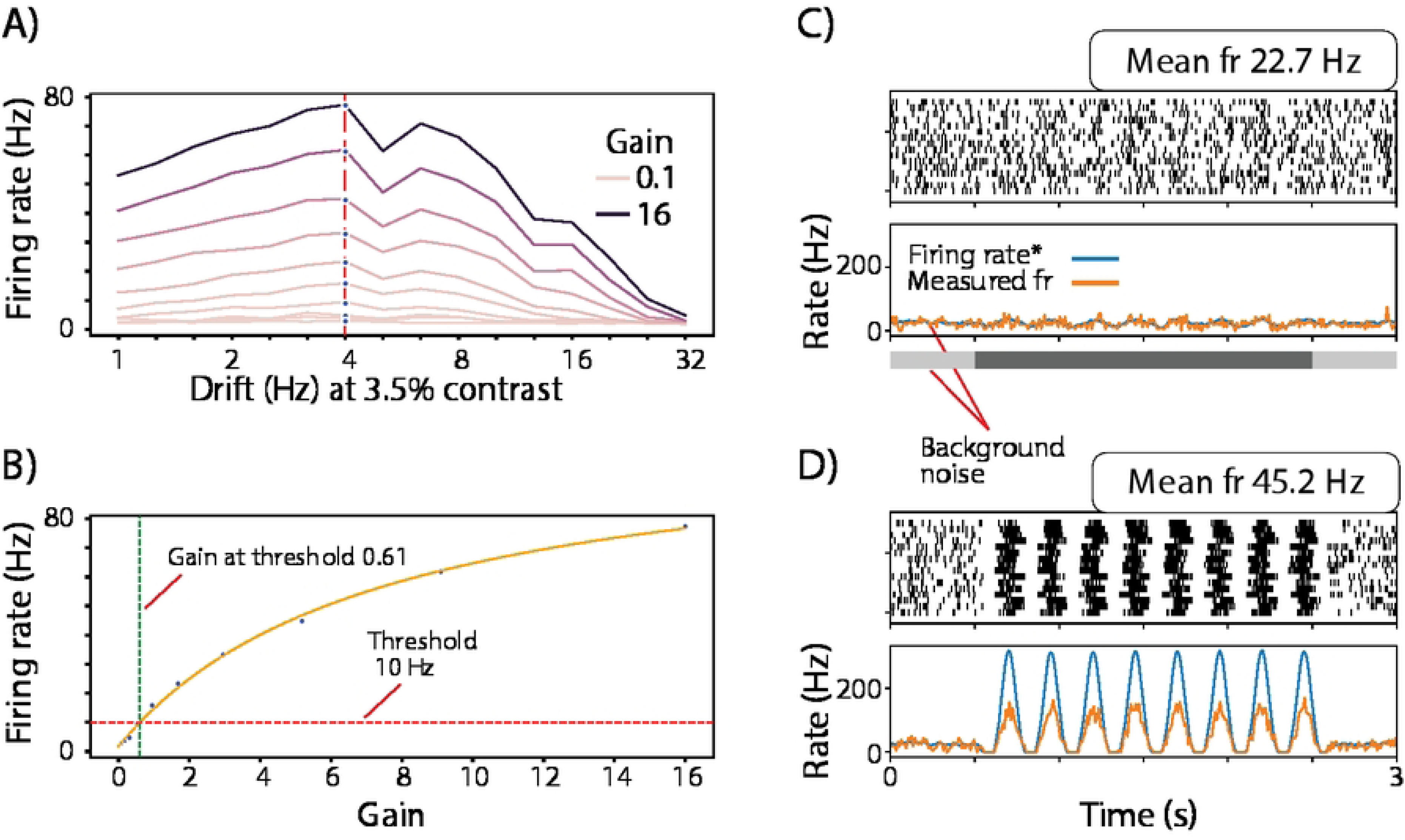
Generator potential gain calibration for parasol ON DOG Fixed model. A) Multiple gain values were simulated across a range of grating drift frequencies. After Fourier transform, the frequency with the largest number of peak values was selected for reference (red dashed line with blue dots). B) The response for the reference drift frequency increase nonlinearly. Orange solid line: descriptive Naka-Rushton function. Green dashed line: Gain producing the artificial “detection threshold” of 10 Hz, marked with the red dashed line. This gain value became the system gain for the parasol ON DOG fixed model combination. The same procedure was repeated for the other model combinations C) The parasol ON unit responses for 2-s drifting grating stimulus at the threshold contrast (3.5%). Blue curve: requested firing rate (Firing rate*). Orange curve: measured firing rate. D) As in C) but for 100% contrast.

Figure 6C shows the parasol ON unit responses for 2-s drifting grating stimulus at the threshold contrast (3.5%). The signal is dominated by background noise, and the requested firing rate (blue curve) and measured firing rates (orange curve) are similar. For 100% contrast (Fig 6D), the actual spike rate is limited by the refractory spike generation model. A Poisson model would be aligned with the requested firing rate (not shown).

### 2.8 Software implementation

The software was written in Python and uses the unit and dynamic equation systems of Brian2 [57]. A GPU acceleration option is flexibly available both via Brian2Cuda [58] for Dynamic temporal model and Pytorch (https://pytorch.org/) for VAE model training and multiple general computing tasks. Development and standalone simulations were run on a laptop with 64GB RAM and an NVIDIA GeForce RTX 3080 Ti GPU. The main results involving several hundred simulations (gain calibrations and Figures 8, 9, and 10) were computed on the University of Helsinki computing cluster or on the national IT Center for Science (CSC) supercomputer Puhti using a containerized environment.

## 3 Results

Figure 7 shows the model components and optional pathways from stimulus video to spike generation. Below, we present a set of model combinations showing a good match to experimental data, as well as the comparison to a subset of other model combinations. In Figures 8, 9 and 10 the line color indicates the experimental data (black), temporal model type (blue, orange, green) or control simulation data (magenta). The thick line indicates parasol and thin line midget responses.

**Figure 7.**
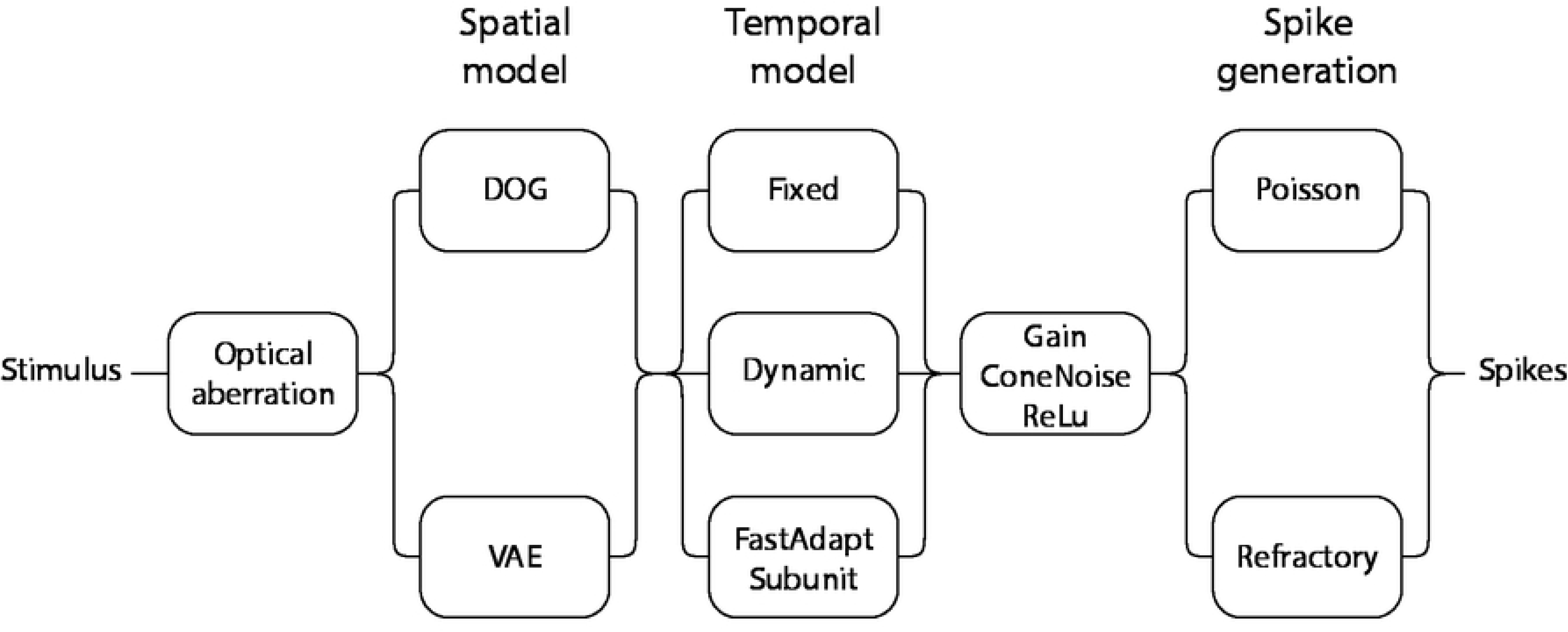
Overview of modeling process and the spatial, temporal, and spike generation model variants.

**Figure 8.**
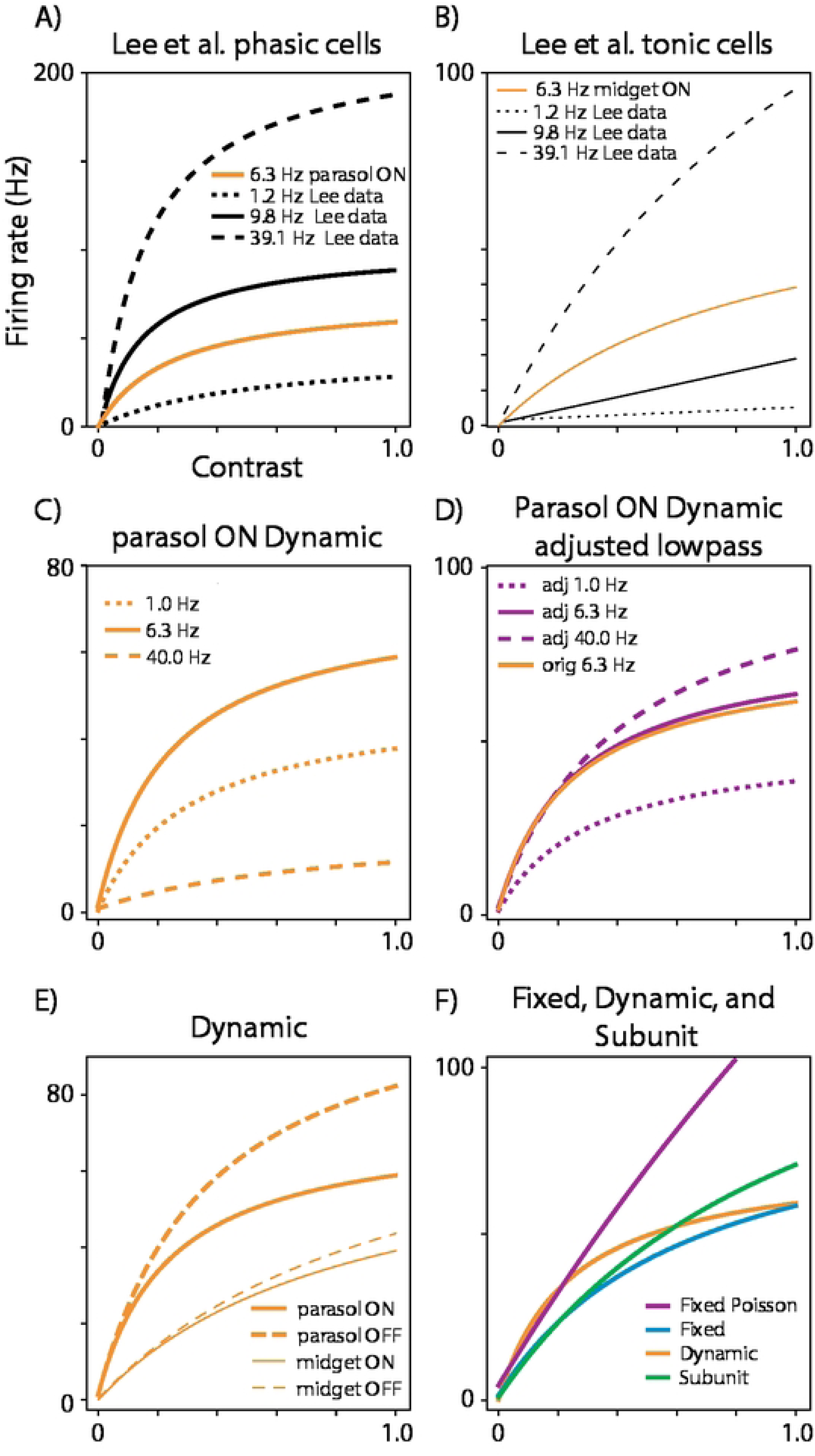
Contrast response function to full-field flicker stimulus. The data in A) and B) are digitized from Lee et al., [59]. The data was fit with Naka-Rushton function [88]: R(c) = baseline + (R_max_ c) / (c_50_ + c), where R(c) is the response, c is the contrast, R_max_ is the maximum response, c_50_ is the contrast at which the response is half of R_max_. The orange lines show simulated Dynamical temporal model responses. C) Contrast response function at difference flicker rates. D) Corresponding flicker rates after increasing the low-pass corner frequency parameter in the Dynamic temporal model. E) Comparison of parasol and midget ON and OFF responses for the Dynamic model. F) Comparison between temporal model types. All simulations use DOG spatial model and include the background activation emerging from the cone noise. The magenta line in F) uses Poisson spike generation mode, whereas all other simulations use the refractory model. The line color indicates the experimental data (black), temporal model type (blue, orange, green) or control simulation data (magenta). The thick line indicates parasol and thin line midget responses.

**Figure 9.**
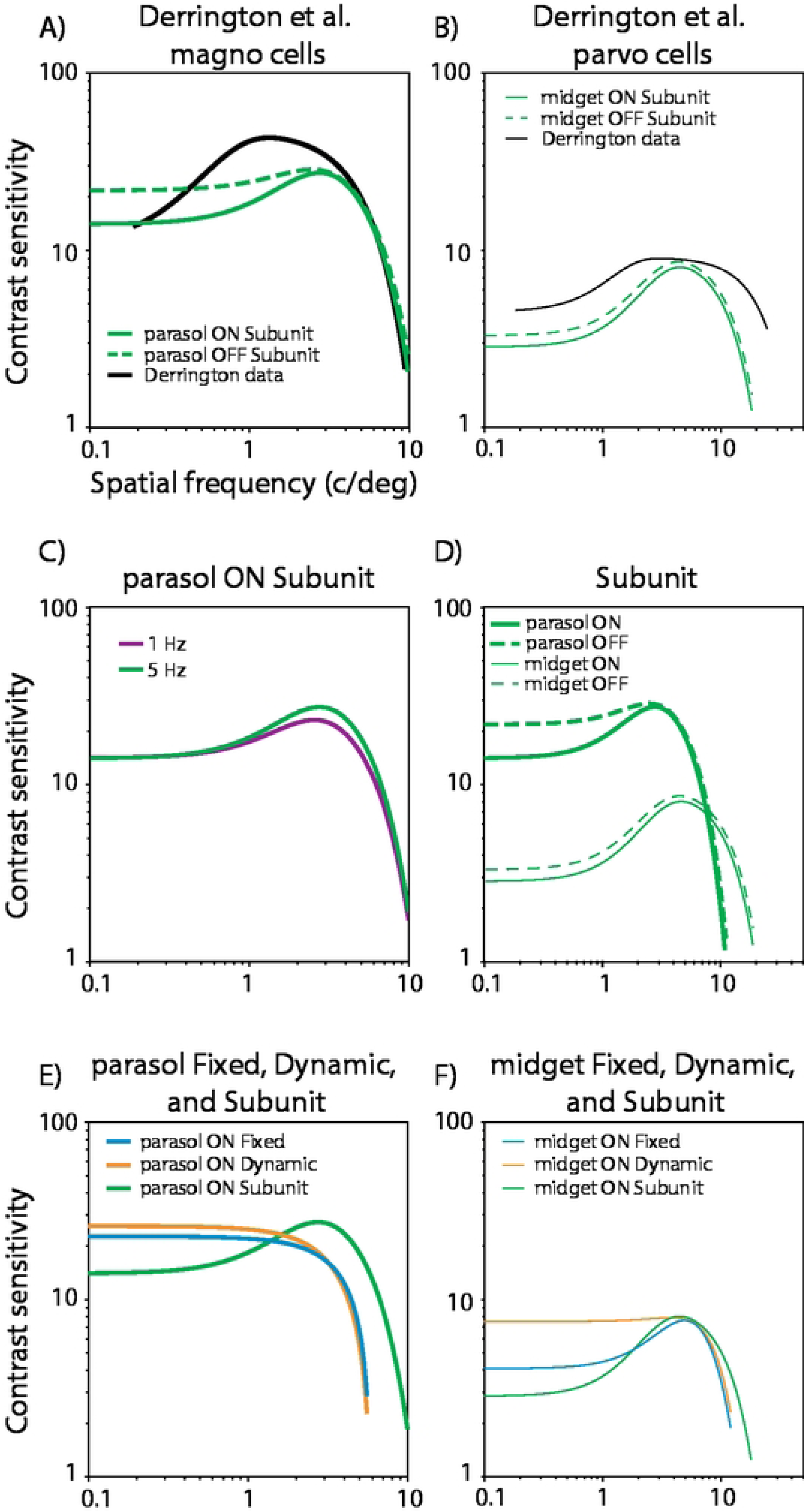
Spatial contrast sensitivity function. Responses were measured with 5-Hz drift frequency. The spatial contrast sensitivity function is calculated as F(sf) = K (k_c_ pi r_c_^2^ exp(-(pi sf r_c_)^2^) - k_s_ pi r ^2^ exp(-(pi sf r_s_)^2^)), where K is the overall sensitivity, k_c_ and r_c_ are the magnitude and radius for the center, and k_s_ and r_s_ are the magnitude and radius for the surround, respectively, from [89]. A single contrast sensitivity function curve represents the mean over units. Parasol (A) and midget (B) data (black curves, from [56]) and Subunit temporal model simulations (green). Spatial contrast sensitivity was estimated in a two-step process. First, varying contrast responses were fitted with the Naka-Rushton model, separately for each spatial frequency. The contrast sensitivity was the reciprocal of the contrast value where the unit responses reached 10-Hz. C) Control experiment with 1-Hz drift frequency. D) Comparison of parasol and midget ON and OFF responses for the Subunit model. Comparison between temporal model types for the parasol (E) and midget (F) units.

**Figure 10.**
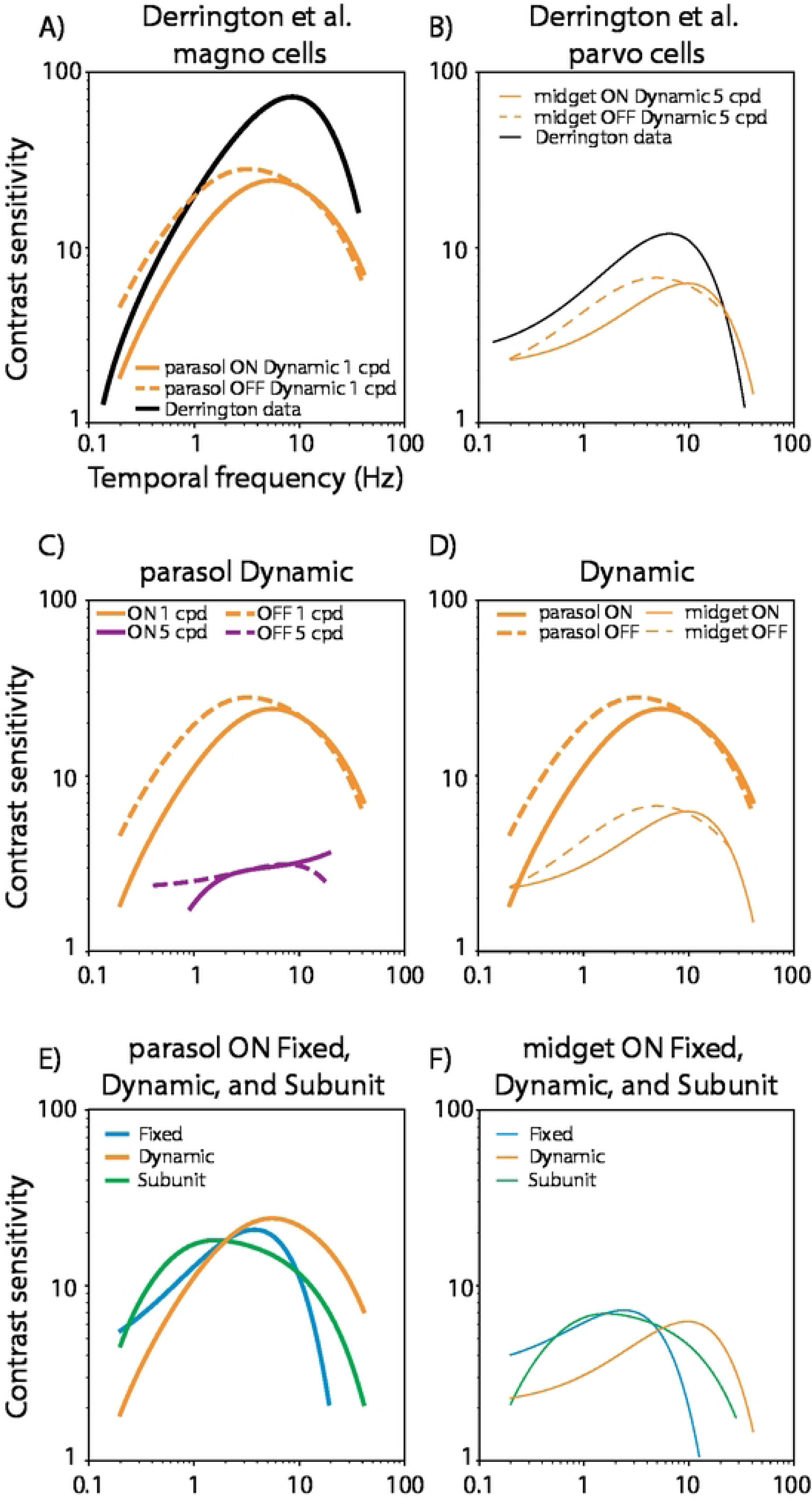
Temporal contrast sensitivity function. Following [56], the temporal contrast sensitivity function is expressed as the difference between two exponentials: F(tf) = s_1_ exp(-tf k_1_) - s_2_ exp(-tf k_2_), where F(tf) is the temporal contrast sensitivity, s_1_ and s_2_ are the scaling constants for the first and second processes, and k_1_ and k_2_ are the reciprocals of the time constants for the first and second processes, respectively. A) Data (black line, from [56]) and Dynamic temporal model simulations for ON (orange solid line) and OFF (orange dashed line) response types with 1 cpd spatial frequency drifting grating. B) Corresponding data and simulations for the midget units with 5 cpd spatial frequency drifting grating. C) Control experiment with parasol units at the 5cpd spatial frequency. D) Comparison of parasol and midget ON and OFF responses for the Dynamic model. Comparison between temporal model types for the parasol (E) and midget (F) units.

### 3.1 Luminance contrast response function

Figure 8 (A, B) shows parafoveal (3-10 deg eccentricity) contrast response function for phasic and tonic cells. Data were digitized from [59], and originally obtained from in vivo recordings of anesthetized macaques. The stimulus was a sinusoidally-modulated full-field luminance flicker. The phasic cells represent the magnocellular retinocortical pathway, originating from parasol ganglion cells, and the tonic cells represent the parvocellular pathway, originating mainly from midget ganglion cells. The phasic cells display the characteristic saturating contrast response function, whereas the tonic cells are more linear. The semi-saturating magnocellular responses and the near-linearity of the parvocellular responses are well-established for both ganglion cells [59,60] and LGN [61,62].

The Dynamic temporal model (Fig 8 A, orange curve) best follows the experimental data and shows a saturating contrast response function for the parasol units and near-linear contrast response for the midget units (Fig 8 B).

Our Dynamic temporal model cannot follow the 40-Hz flicker frequencies (Fig 8 C), as the Lee et al data, suggesting lower low-pass filter cutoff frequency in the experimental parametrization [46]. When we artificially reduce the filter cutoff by reducing the time constant TL (Eq. 4, Fig 8 D) we recover much of the sensitivity of the Dynamic model to the 40-Hz flicker.

For both midget and parasol units, the ON and OFF responses follow similar contrast response functions (Fig 8 E). In contrast to the Dynamic model, the Fixed and Subunit models show less saturation in their contrast response function (Fig 8 F). This minor saturation emerges from the refractory spike generation model, precluding the highest firing rates. With a Poisson spike generation model (Fig 8 F, magenta) the contrast response function becomes linear, as expected from the linearly summing Fixed model.

### 3.2 Spatial contrast sensitivity

Behavioral spatial contrast detection threshold varies with spatial frequency in both humans and macaques [63]. Both species show peak sensitivity at about 5 cpd in low photopic range, and the peak shifts towards lower frequencies for lower background luminance, eventually showing only high-pass with cutoff at about 1 cpd. When a corresponding sensitivity is measured from the LGN, the function shows less low-frequency attenuation than in the behavioral data [56,64].

Figure 9 (A, B, black curves) shows spatial contrast sensitivity functions for the magno- and parvocellular pathways, obtained from the LGN in anesthetized macaques. The magnocellular data was digitized from two cells and the parvocellular data from six cells, both from [56].

The Subunit temporal model (Fig. 9 A, B, green curves) shows the best match to these data. It shows partial low-frequency attenuation, as the experimental data, but the sensitivity peaks for parasol units land above the experimental data.

Figure 9C demonstrates how the responses show little sensitivity to shifts in the temporal frequency of the drifting grating.

Figure 9D shows that the parasol model units show higher sensitivity and peak at lower spatial frequencies, whereas the midget units reach higher spatial frequencies, as in macaque data (Fig. 9 A, B black curves), and expected from their different RF sizes. In addition, the OFF units are consistently somewhat more sensitive than the ON units.

Figure 9 E and F compare the Subunit temporal model to the Fixed and Dynamic models. The characteristic low-pass attenuation is weak for the Fixed and Dynamic models, and the parasol Subunit model is sensitive for somewhat higher spatial frequencies, as expected due to the enhanced resolution at the center. This enhanced resolution emerge from the unit-wise bipolar non-linearity, which we modelled following [24].

### 3.3 Temporal contrast sensitivity

Figure 10 A and B show contrast sensitivity as a function of varying grating drift frequency. The experimental data is from the LGN and the black curves show a double exponential function fit to data from two cells each [56]. Both parasol and midget Dynamic temporal model simulations show similar temporal sensitivity profile to experimental data.

The parasol units at 5 cpd spatial frequency (Fig 10C) clearly lose sensitivity to the stimulus.

Figure 10D compares the midget and parasol units from A and B panels. While parasol units have stronger low-frequency attenuation, the high-frequency cutoff is similar between the unit types, like in experimental data (Fig. 10 A, B black curves).

Figures 10 E and F compare the Fixed, Dynamic and Subunit temporal models for parasol and midget units. All models show temporal band-bass character, but the sensitivity peaks house distinct frequencies, the Dynamic being closest to experimental data (Fig. 10 A, B).

### 3.4 Natural video response reveals the distinct population dynamics for the unit types

Figure 11 shows the DOG spatial and subunit temporal model responses for all cell types for a head-mounted natural video input [65]. The Midget OFF unit spike rate emphasizes the local dark contrasts (overlaid onto video frames). The Midget OFF unit rasterplot has ∼5-Hz noise before stimulus onset at 0.5-s. The Midget ON unit has a ∼25-Hz noise background, and a complementary activation pattern to Midget OFF units. These distinct noise levels are estimates from macaque retina recordings (Raunak Sinha, personal communication). The Parasol ON and OFF units show similar complementary activation patterns, but a higher mean spike rate. The bottom two rasterplots compare the Subunit and Fixed temporal model responses. While both models were calibrated to give the same spike rate for low-contrast gratings, the Subunit model is more responsive than the Fixed model for the natural video.

**Figure 11.**
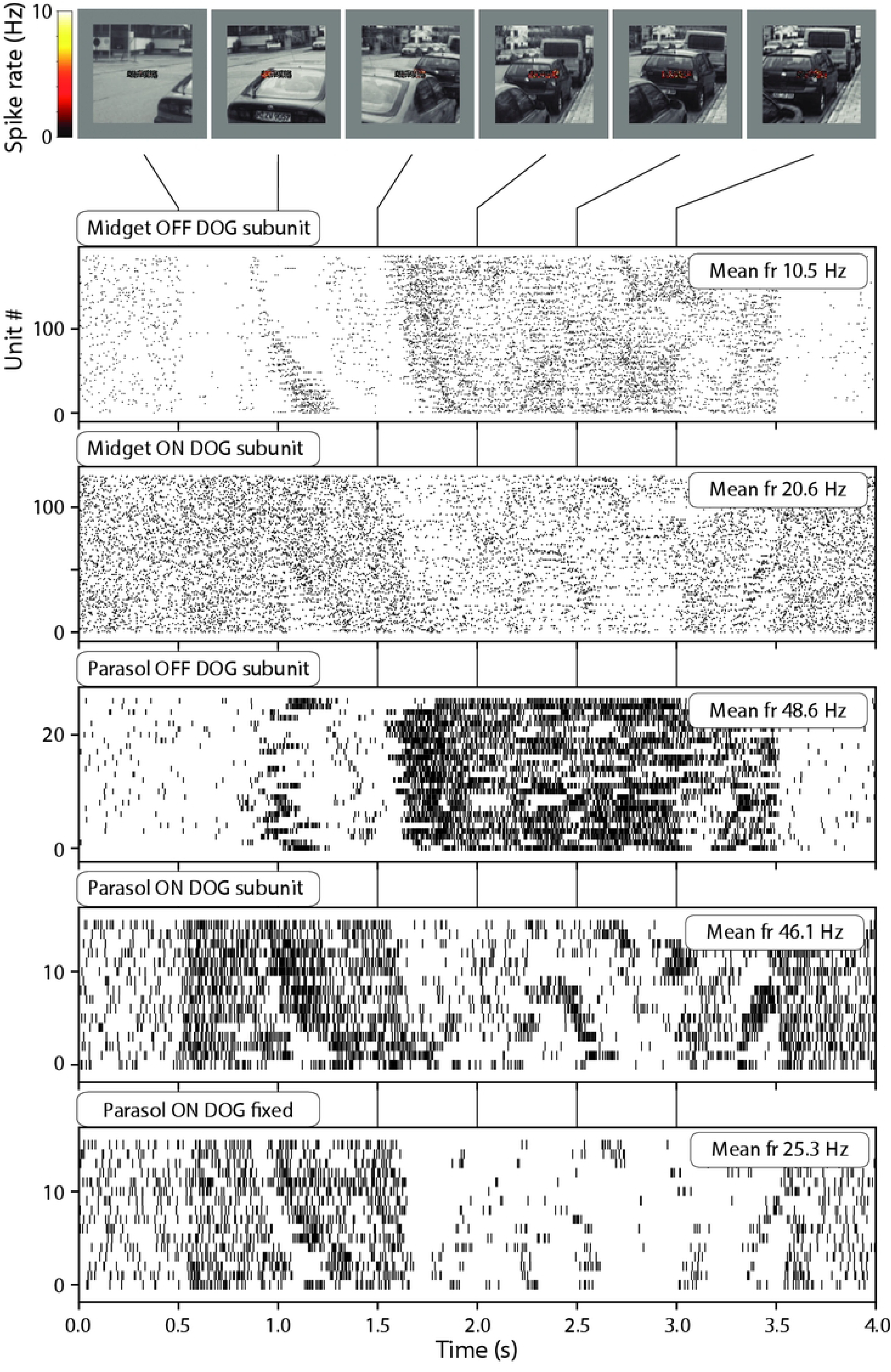
Retina response to natural head-mounted video input (from [65]). Top. A video stimulus with 0.5 second pre- and poststimulus gray baseline. Midget OFF unit positions and spike rates (100-ms sliding average) are overlaid onto the video frames. Bottom. Spike raster plots for the video stimulus for different ganglion cell types.

### 3.5 A shared cone noise model preserves spectral properties and temporal dependency between units at short distances

Primate retinal ganglion cells share cone noise locally across cell types and this cone noise is the primary driver of ganglion cell background activation [27]. The downstream processes are limited by this noise, but in addition they might utilize such noise spike synchrony e.g. for associating spatially overlapping receptive fields during development and maintenance of spatial tuning.

Figure 12 examines the temporal and spatial correlation of noise for Parasol and Midget ON units; both unit types occupy the same retina patch (Figure 12 A). Figure 12B shows the cone noise data and model [53], together with the shared Parasol ON firing rate spectral response (blue line). The model output attenuates at the high frequencies, following the cone dynamics, albeit somewhat low-pass filtered. The Gaussian model (independent noise, red line) loses the temporal dependencies, resulting in whitened spectral response. Figure 12C shows that some units show temporal correlations which are lost for the Gaussian noise model. Correlations emerge both within (top) and between (bottom) unit types. Figure 12D shows that the temporal dependencies are spatially local in the retina model.

**Figure 12.**
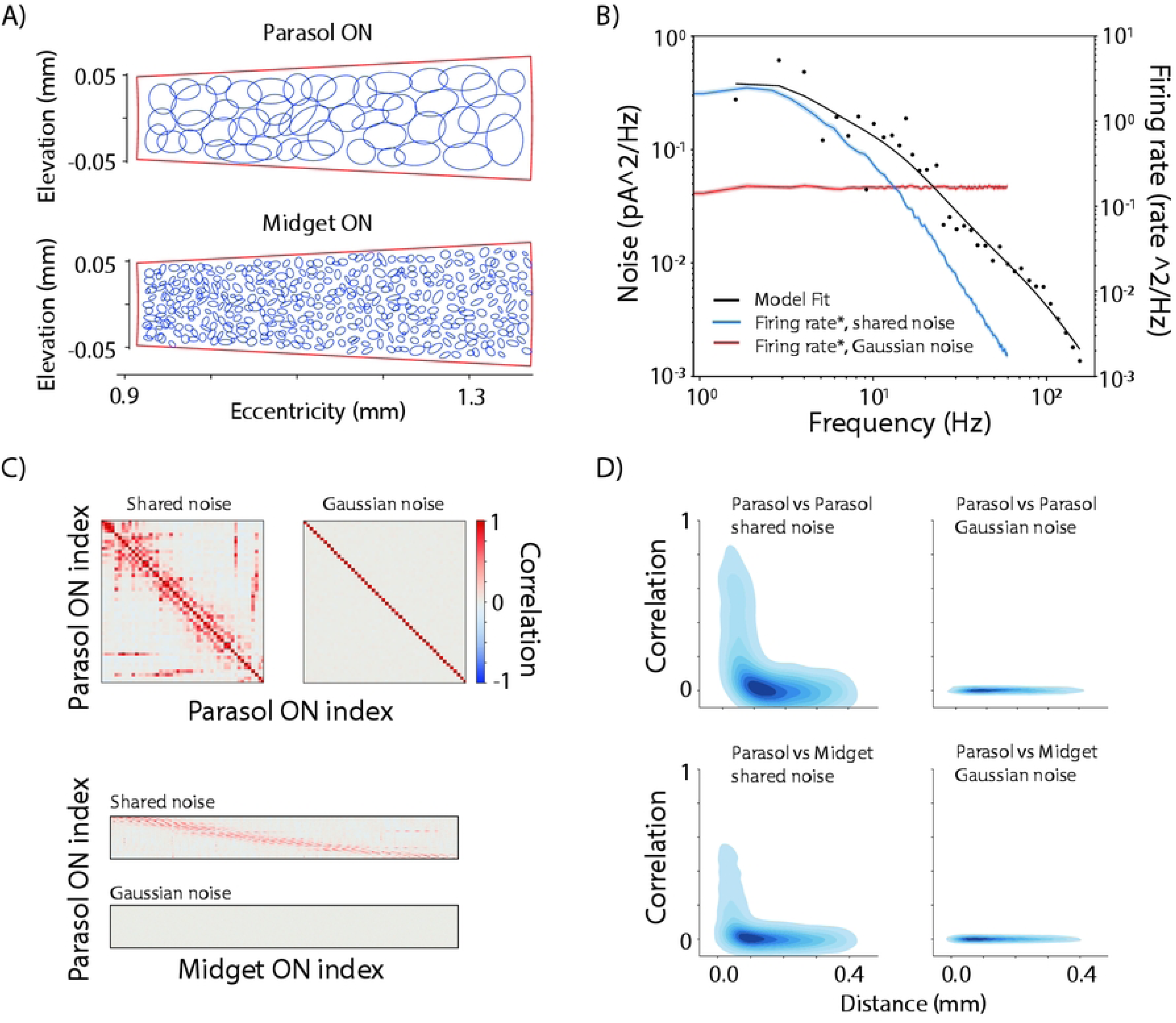
Noise characteristics. A) Parasol and midget ganglion cell mosaics centered at 5 deg eccentricity along the horizontal meridian. Reduced density (80% of literature values) for clarity. B) Black dots: cone noise spectral data from [53]. Black line: A fitted model composed of scaled cone response (at 4k R*), and two Lorenzian functions (at 24 and 200 Hz corner frequencies). Blue line and shading: parasol unit requested firing rate (before spike transition, denoted with *) for shared and Gaussian noise models. C) Firing rate* correlation between units. Top: Parasol ON unit correlations. Bottom: Parasol ON and Midget ON unit correlations. D) Kernel density estimate -plots for the distance dependence of correlations.

## 4 Discussion

In this paper, we summarized anatomical and physiological data and implemented a set of models of macaque retina with the aim of providing biologically plausible input for simulations of the visual cortex. Our macaqueretina Python package can construct artificial patches of retina with ON and OFF midget and parasol populations, with each cell type producing spiking responses for video input. Apart from gain calibration at threshold contrast, the parameters were fixed to distributions extracted from literature and earlier data. We validated these models against well-established measurements of visual sensitivity, namely luminance flicker contrast response functions, as well as spatial and temporal contrast sensitivity functions.

### 4.1 Earlier retina simulation software

Earlier retinal simulators provide a detailed account on optics, eye movements and cone signals [66], emphasize biophysical unit dynamics [68], model percept from retina prosthesis [69], or provide simple spiking retina models emphasizing real-time processing [70], but these are not applicable for visual cortex simulations which need larger retinal patches providing a substantial proportion of ganglion cell spiking output. There are some tools for simulating on larger patches of the retina [71–74]. The CONVIS simulator supports complex spatiotemporal receptive fields, including V1 responses [71]. However, it remains unclear how one should parametrize them to obtain models for the main macaque ganglion cell types. In addition, eccentricity-variant sampling options are limited, which is an essential feature of the primate retina and retinocortical mapping. Third, no simulator includes correlated noise with natural spectral characteristics which in our case is the main driver of the background ganglion cell response variability [27].

### 4.2 Limitations of the macaque retina simulator

We cannot provide one comprehensive, or “true”, macaque retina model, because such model does not yet exist. Instead, we provide a combination of spatial, temporal and spike generation models which are apt for distinct visual cortex simulation experiments. For a detailed spatial feature space, a combination of VAE spatial model with Subunit temporal model is most appropriate, whereas for learning dynamical features from the input likely the Dynamic temporal model with refractory spike generation is more appropriate.

The division of vision into P and M pathways driven by midget and parasol GCs is obviously a simplification, as a growing a number of distinct GC types have been identified as projecting to the LGN [39,75]. Still, the overwhelming majority of input to the parvo- and magnocellular layers comes from midget and parasol cells, respectively, while the newly discovered GC types each constitute only 1-2% of GCs [2]. Low proportion, however, does not imply negligible functional significance of these groups for photopic image-forming vision.

The scope of our set of models is limited in several ways. First, we focused on the macaque retina but presumably the models can be reparametrized for other primate species, including humans. For rodents, such reparameterization has limited value because of their qualitatively distinct retinal processing [1]. Second, the model is restricted to achromatic stimuli; adding chromatic sensitivity requires at least one additional ganglion cell type and implementation of the chromatic cone primaries upstream, as in ISETBio [66]. Third, we do not model the whole retina but the temporal hemiretina; the nasal hemiretina comprises distinct cell density profile and the optic nerve head close to the horizontal meridian.

The early visual pathway is challenged by the variability of statistical properties in natural scenes. For example, contrast and luminance vary 10-fold in typical natural scenes [76]. Thus, rapid contrast and luminance gain control is needed. In response to luminance decrease, both parasol and midget GCs change their temporal tuning to prefer lower temporal frequencies and their tuning curves become more low-pass in shape [59,77]. With fixed mean luminance, only the parasol GCs significantly adapt their gain and integration time in response to changes in contrast [60,78–80] which we aim to follow with our Dynamic temporal model. Our Subunit model includes fast cone adaptations whose dynamics change with background illumination, but here we clearly miss the fastest adaptation (Fig 5B).

Adaptation to luminance and contrast seems to operate independently from each other [81]. We have omitted the slow form of contrast adaptation in parasol GCs that reduces gain without affecting the temporal response [82,83].

For the spatial RF, increase in contrast has been shown to increase the strength of the RF surround in cats and New World monkeys [84,85]. In line with these nonlinearities, an artificial recurrent neural network performs better than a generalized linear model, even with added scalar nonlinearities, in predicting the spiking of parasol GCs in response to stimulation using natural images [86]. We have omitted this contrast dependence of the spatial surround strength.

Finally, there remains a large gap between any heuristics in retinocortical pathway development for modelling purposes and the actual biological development [87]. When functionally relevant retinal features become more comprehensively characterized in the future, we are likely to have more compute efficient as well as natural retinal spike generators. One option is to fit a temporospatial nonlinear filter to large set of data, which might be feasible for the CONVIS simulator [71].

### 4.3 Future prospects

Our macaqueretina simulator will require further development and validation depending on the scope of cortical simulation studies. As such it has been tested with artificial stimuli against existing old macaque data. With proper data, it should be straightforward to reparametrize the models for other primate species or validate further for natural images and videos. We hope that this open source and modular software will encourage not only reparameterizations but also modifications and extensions for related research questions.

## Financial Disclosure Statement

This work was supported by Research Council of Finland grant N:o 361816 to SV. The funders had no role in study design, data collection and analysis, decision to publish, or preparation of the manuscript.

## Data availability statement

There are no primary data in the paper; we have archived our code and included statistics to create new retina models on Zenodo (DOI <u>10.5281/zenodo.17726225</u>). Modifiable parameters are available inside the repository in human readable format (yaml files). The code is released under the MIT license.

## Acknowledgments

We thank E.J. Chichilnisky who kindly allowed us to access their spatiotemporal receptive field data, James Golden for help with interpreting the receptive field data. We thank Petri Ala-Laurila for insightful comment on model construction, Raunak Sinha for providing estimates of ganglion cell background firing rates and Viljami Salmela for help with the natural video stimuli. We thank Silviya Hasana for pointing out the Derrington 1984 work, which we use as validation data.

## Author contributions

HH conceived the study, programmed the software and wrote the manuscript.

SV conceived the study, programmed the software and wrote the manuscript.

FV programmed the software and wrote the manuscript.

## Author information

Competing financial interests: None.

## Notes

### Competing Interest Statement

The authors have declared no competing interest.

